# Effects of hydroxychloroquine on atrial electrophysiology in *in silico* wild-type and PITX2^+/−^ atrial cardiomyocytes

**DOI:** 10.1101/2022.12.07.519115

**Authors:** Euijun Song

## Abstract

Hydroxychloroquine (HCQ) is commonly used in the treatment of autoimmune diseases and increases the risk of QT interval prolongation. We quantitatively examined the potential atrial arrhythmogenic effects of HCQ on atrial fibrillation (AF) using a computational model of human atrial cardiomyocytes. We measured atrial electrophysiological markers after systematically varying HCQ concentrations. HCQ concentrations were positively correlated with the action potential duration (APD), resting membrane potential, refractory period, APD alternans threshold, and calcium transient alternans threshold (P<0.05). In contrast, HCQ concentrations were inversely correlated with the maximum upstroke velocity and calcium transient amplitude (P<0.05). When the therapeutic concentration (C_max_) of HCQ was applied, HCQ increased APD_90_ by 1.4% in normal sinus rhythm, 1.8% in wild-type AF, and 2.6% in PITX2^+/−^ AF, but did not affect the alternans thresholds. The overall *in silico* results suggest no significant atrial arrhythmogenic effects of HCQ at C_max_, rather implying a potential antiarrhythmic role of low-dose HCQ in AF. However, at HCQ of 4-fold C_max_, a rapid pacing rate of 4 Hz could induce prominent APD alternans, particularly in the PITX2^+/−^ AF model. Concomitant PITX2 mutations and HCQ treatments may increase the risk of AF, and this potential arrhythmogenic effect of HCQ should be further investigated.

## 1. Introduction

Hydroxychloroquine (HCQ) is an antimalarial drug commonly used to treat certain types of autoimmune diseases, such as systemic lupus erythematosus (SLE), rheumatoid arthritis, and Sjogren’s syndrome [1, 2]. HCQ has also been tested for treatments and prophylaxis of COVID-19 but did not show significant efficacy in treating/preventing COVID-19 [3–5]. Various mechanisms of action of HCQ have been discovered [2]. HCQ interferes with Toll-like receptor (TLR) signaling pathways and inhibits lysosomal activity and autophagy. In the immune system, HCQ reduces the production of tumor necrosis factor (TNF), interleukin (IL)-6, and interferon (IFN)α in plasmacytoid dendritic cells, thereby inhibiting the type-I IFN signaling in SLE [6–8].

Based on multiple immunomodulatory actions, SLE is primarily managed with HCQ to reduce disease activity, flare risk, and organ damage [9]. However, long-term and/or high-dose HCQ treatments can lead to several adverse effects, including QT prolongation, cardiomyopathy, and retinopathy [10–12]. Specifically, HCQ-induced QT prolongation is directly linked to life-threatening ventricular arrhythmias, such as *Torsade de Pointes* (TdP) [13, 14].

HCQ inhibits several human cardiac ion channels, including rapidly activating delayed rectifier potassium current (I_Kr_), inward rectifier potassium current (I_K1_), fast sodium current (I_Na_), and L-type calcium current (I_CaL_) channels [15, 16]. The net effects of HCQ on cardiac ion channels can prolong action potential duration (APD) in ventricular cells [15–17], eventually leading to QT interval prolongation. To examine the potential proarrhythmogenic risk of non-cardiovascular drugs such as HCQ, computational modeling-based drug toxicity screening frameworks (e.g., CiPA [18, 19]) have been proposed and tested [15, 16]. These computational electrophysiology models have been developed based on quantitative patch-clamp data of human cardiomyocytes [20, 21] and recently utilized to perform computational modeling-guided ablation of atrial fibrillation (AF) [22, 23] or *in-silico* antiarrhythmic drug trials [24, 25]. However, most recent preclinical studies have focused on the HCQ-associated TdP risk stratification in ventricular cells. It is still unclear how HCQ affects atrial electrophysiology and the risk of AF.

In a retrospective cohort of 1,647 adult patients with SLE, Gupta et al. [26] discovered that HCQ use was associated with an 88% decreased risk of new-onset AF. This AF-protective effect of HCQ remained consistent in the propensity score-matched analysis. These results suggest that HCQ may play an antiarrhythmic role in AF in patients with SLE. However, potential antiarrhythmic mechanisms of HCQ have remained unknown, and further prospective clinical study is needed in a general population.

In this computational study, we systematically examine the potential effects of HCQ on atrial electrophysiology using computational models of human atrial cardiomyocytes. We quantitatively analyze HCQ-induced changes in APD, calcium transient (CaT), and APD/CaT alternans thresholds by varying HCQ doses in normal sinus rhythm (NSR) and AF conditions. Since PITX2 plays a key role in AF electrical/structural remodeling and genetic variants located on 4q25, an intergenic region near the *PITX2* gene, are closely associated with AF [27–30], we also examine the effects of HCQ in an *in silico* PITX2^+/−^ AF model.

## 2. Methods

### 2.1 Computational modeling of human atrial cardiomyocytes

We used a state-of-the-art mathematical model of the human atrial cardiomyocyte developed by Grandi et al [20]. The Grandi atrial cell model incorporates comprehensive Ca^2+^ handling properties in both normal and AF conditions. Briefly, the cell model is described according to the following differential equation:

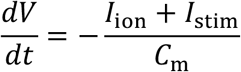

where *V* is the transmembrane potential, I_ion_ is the total ionic current, I_stim_ is the stimulus current, and C_m_ is the total membrane capacitance. Each ionic current is a variation of the Hodgkin-Huxley-type model, that is, a nonlinear function of the action potential, ion concentration, and ion channel gating state variables. The biophysical details are described in the study by Grandi et al [20]. Numerical simulations were performed with MATLAB 2021b, and ordinary differential equations were numerically solved using a variable-step, variable-order method (*ode15s* function in MATLAB).

To reflect the electrical remodeling in AF, we decreased I_Na_, I_CaL_, I_to,fast_, I_Kur_, I_SERCA_, increased I_K1_, I_Ks_, I_NCX_, I_RyR_, I_SR,leak_, and added an I_NaL_ component as described in Grandi et al [20]. The PITX2^+/−^ AF condition was modeled by reducing I_K1_ by 25% and doubling I_Kr_ from the wild-type AF baseline condition [27].

### 2.2 Measurement of atrial electrophysiological markers

At a pacing cycling length (PCL) of 1000 ms (=1 Hz), we measured the APD at 90% of repolarization (APD_90_), APD at 50% of repolarization (APD_50_), resting membrane potential (RMP), maximum upstroke velocity (V_max_), and calcium transient amplitude (ΔCaT) [31, 32]. A steady state was achieved by stimulating the cells for 60 s. The ΔCaT was defined as the difference between the maximum and minimum intracellular Ca^2+^ concentrations [31–33]. We also estimated the refractory period and APD (or ΔCaT) alternans threshold cycle lengths using a dynamic ramp pacing protocol as previously described [32]: PCLs of 1000, 600, and 500 ms were used, and the PCL was decreased to 300 ms in steps of 50 ms and further decreased in steps of 10 ms until failure of 1:1 capture. For each PCL, the cell was stimulated for 16 beats. The APD (or ΔCaT) alternans were evaluated by determining the maximum and minimum APD (or ΔCaT) values in the last three beats that differed by more than 2% [32]. We considered the beat-to-beat APD variation>10% as *prominent* APD alternans.

### 2.3 Actions of hydroxychloroquine on cardiac ion channels

A drug block was modeled by reducing ionic current conductance values [34]. The ionic current conductance was scaled using the Hill equation as follows:

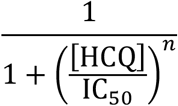

where [HCQ] is the HCQ concentration, IC_50_ is the half-maximal inhibitory concentration, and *n* is the Hill coefficient. As shown in Table 1, we used HCQ ion channel binding profiles in the study by Thomet et al. [15] and assumed the Hill coefficient of 1 [34]. The maximum free therapeutic plasma concentration (C_max_) of HCQ was 0.495 μM (215 ng/mL) [35]. Since HCQ has a long half-life of approximately 40 days, patients with SLE taking daily HCQ doses have blood HCQ concentrations of 1,000–2,000 ng/mL [36]. Thus, we systemically changed the HCQ concentrations from 0.5×C_max_ to 16×C_max_ in a log2 scale (i.e., 0, 0.5, 1, 2, 4, 8, 16×C_max_).

**Table 1.** The maximum free therapeutic plasma concentration (C_max_) and half-maximal inhibitory concentration (IC_50_) values of hydroxychloroquine for human cardiac ion channels based on Thomet et al [15].

| | Value ($\mu M$) |
| --- | --- |
| $C_{\max}$ | 0.495 (=215 ng/mL) |
| $IC_{50}$ | |
| $I_{CaL}$ | 90.0 |
| $I_{Kr}$ | 3.42 |
| $I_{K1}$ | 29.28 |
| $I_{Na,peak}$ | 96.2 |
| $I_{Na,late}$ | 64.9 |
| $I_{Ks}^*$ | >100 |
| $I_{to}^*$ | >4000 |

\* $IC_{50}$ >100 $\mu M$ were excluded from the present analysis.

### 2.4 Statistical analysis

All the numerical simulations and computational analyses were performed with MATLAB 2021b. The ordinary differential equations used in our simulation are deterministic (not stochastic); thus, we did not perform repetitions. The HCQ concentration–electrophysiological marker relationships were evaluated using Spearman’s correlation coefficients and visualized in symmetrical log scales using Python 3.8. P<0.05 was considered statistically significant.

## 3. Results

### 3.1 Hydroxychloroquine-induced changes in atrial action potential and calcium transient

We systematically changed the HCQ concentrations from 0.5×C_max_ to 16×C_max_ and measured action potentials and CaTs in *in silico* atrial cardiomyocytes. As shown in Figure 1, HCQ increased the APD and decreased ΔCaT at a pacing frequency of 1 Hz in a dose-dependent manners. The APD prolongation reflects the outward K^+^ current inhibitions by HCQ, reducing phase 3 of the action potentials. The reduction in ΔCaT may be attributed to the inward Ca^2+^ inhibition by HCQ. The results were consistent in the NSR, wild-type AF, and PITX2^+/−^ AF conditions. We also measured action potentials and CaTs at a pacing frequency of 2 Hz (Figure S1). The dose-dependent HCQ effects on the APD and ΔCaT were similar to those acquired at a pacing frequency of 1 Hz.

**Figure 1.**
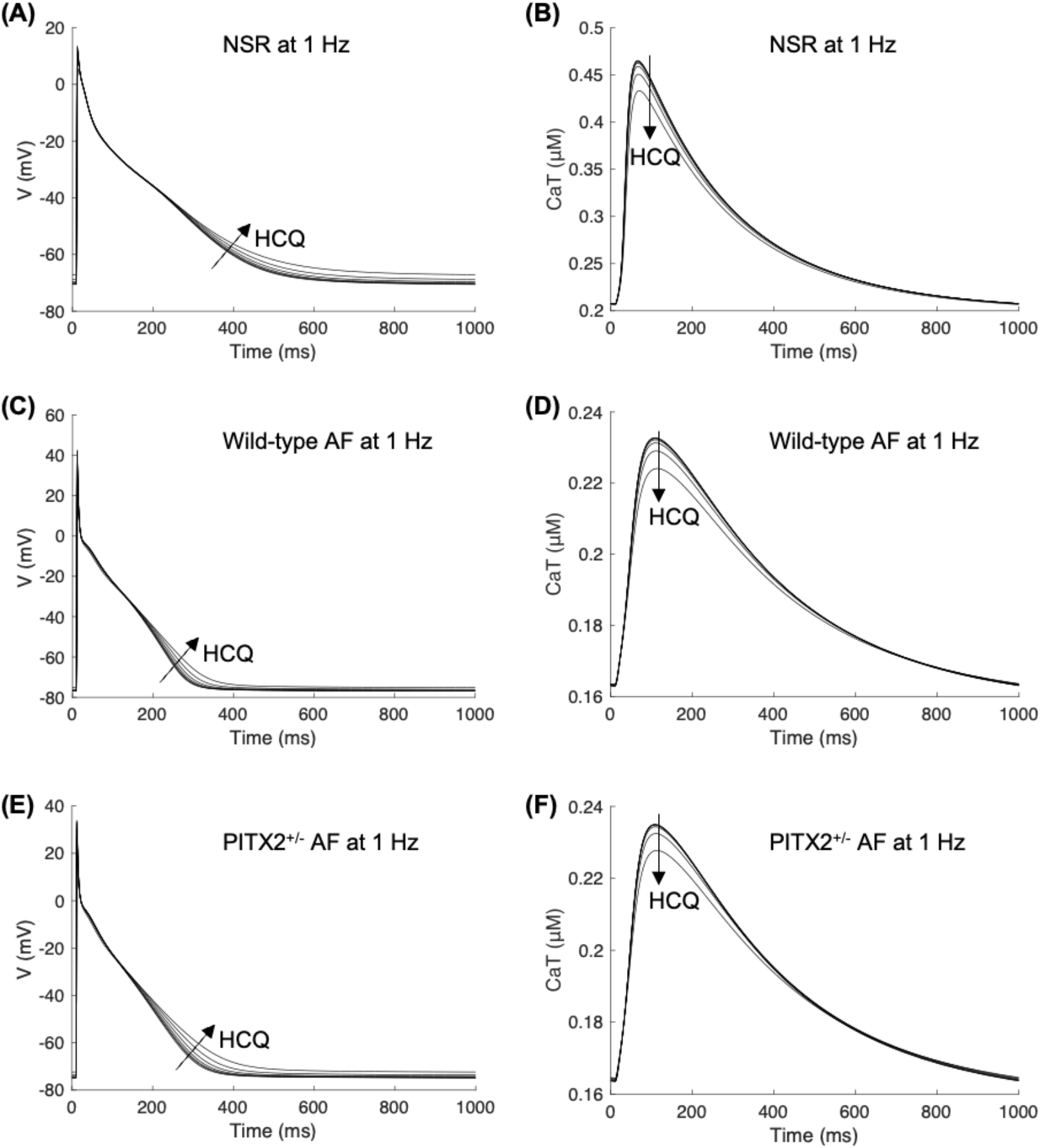
Action potentials (APs) and calcium transients (CaTs) for hydroxychloroquine concentrations of 0–16×C_max_. Simulated steady-state AP and CaT traces are shown at the pacing frequency of 1 Hz in normal sinus rhythm (A, B), wild-type AF (C, D), and PITX2^+/−^ AF (E, F). HCQ, hydroxychloroquine; C_max_, maximum free therapeutic plasma concentration; CaT, calcium transient; NSR, normal sinus rhythm; AF, atrial fibrillation.

### 3.2 Effects of hydroxychloroquine on atrial electrophysiological markers

To quantitatively evaluate the effects of HCQ on atrial electrophysiology, we measured various electrophysiological markers in the NSR (Figure 2), wild-type AF (Figure 3A), and PITX2^+/−^ AF (Figure 3B) conditions. HCQ concentrations were positively correlated with the APD_90_, APD_50_, RMP, refractory period, APD alternans threshold, and ΔCaT alternans threshold (all P<0.05). In contrast, HCQ concentrations were inversely correlated with V_max_ and ΔCaT (all P<0.05). When a therapeutic concentration of HCQ (C_max_) was applied, HCQ increased the APD_90_ by 1.4% in NSR (409.6 to 415.3 ms), 1.8% in wild-type AF (242.8 to 247.1 ms), and 2.6% in PITX2^+/−^ AF (266.8 to 273.8 ms). In contrast, HCQ reduced ΔCaT by 0.5% in NSR (0.258 to 0.257 μM), 0.5% in wild-type AF (0.0699 to 0.0696 μM), and 0.1% in PITX2^+/−^ AF (0.0714 to 0.0714 μM). At C_max_, HCQ did not increase the APD and ΔCaT alternans thresholds in the NSR, wild-type AF, and PITX2^+/−^ AF conditions. Overall, the HCQ-induced changes in atrial electrophysiological markers suggest no significant atrial proarrhythmogenic effects of HCQ at C_max_ (Figures 2 and 3).

**Figure 2.**
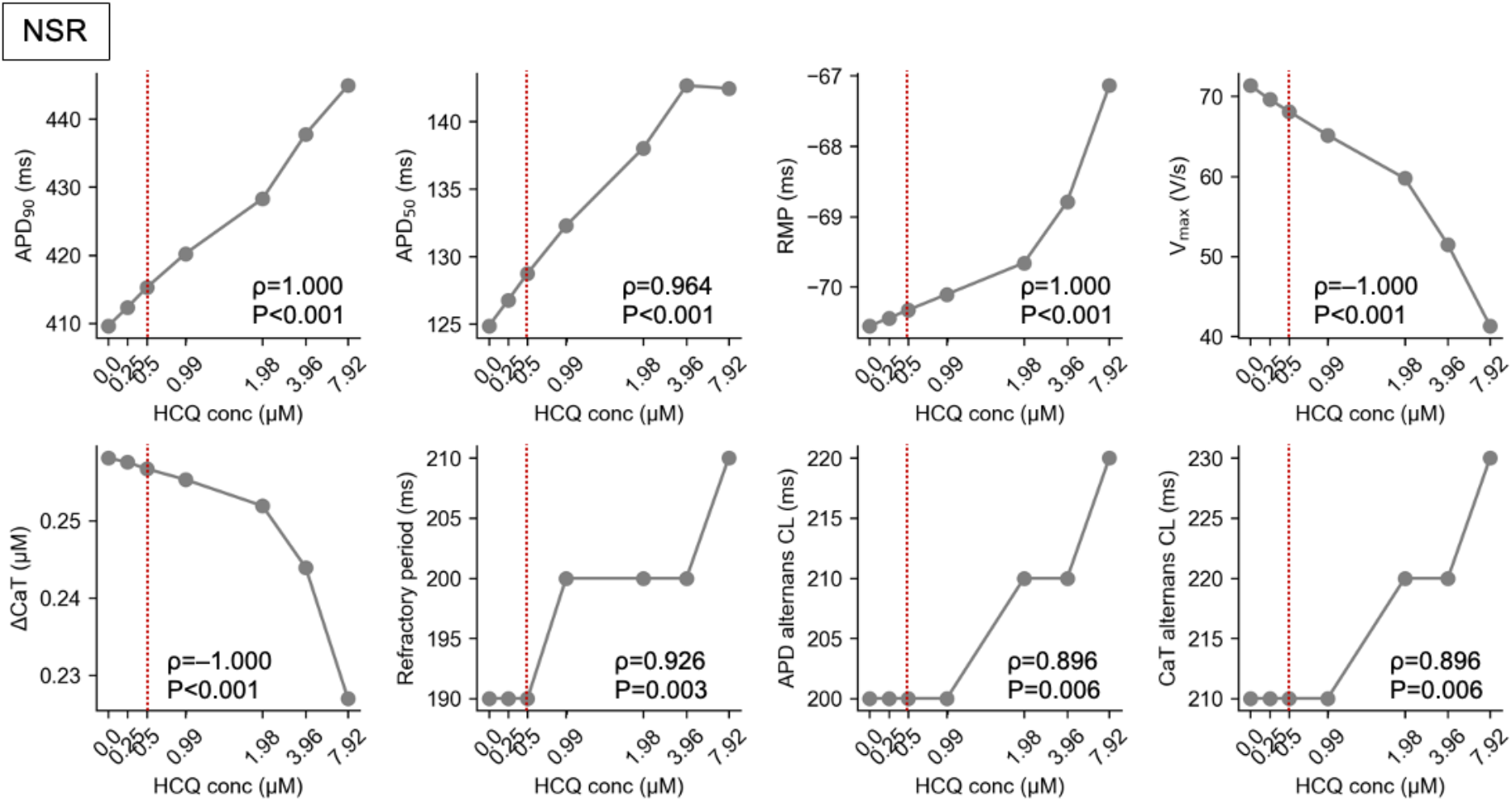
Electrophysiological markers in the normal sinus rhythm (NSR) condition. The action potential duration at 90% of repolarization (APD_90_), APD at 50% of repolarization (APD_50_), resting membrane potential (RMP), maximum upstroke velocity (V_max_), calcium transient amplitude (ΔCaT), refractory period, APD alternans threshold cycle length (CL), and CaT alternans threshold CL are shown, depending on hydroxychloroquine concentrations of 0–16×C_max_ (see Methods). Red lines indicate C_max_. Spearman’s correlation coefficients and P-values are presented. HCQ, hydroxychloroquine; C_max_, maximum free therapeutic plasma concentration; NSR, normal sinus rhythm.

**Figure 3.**
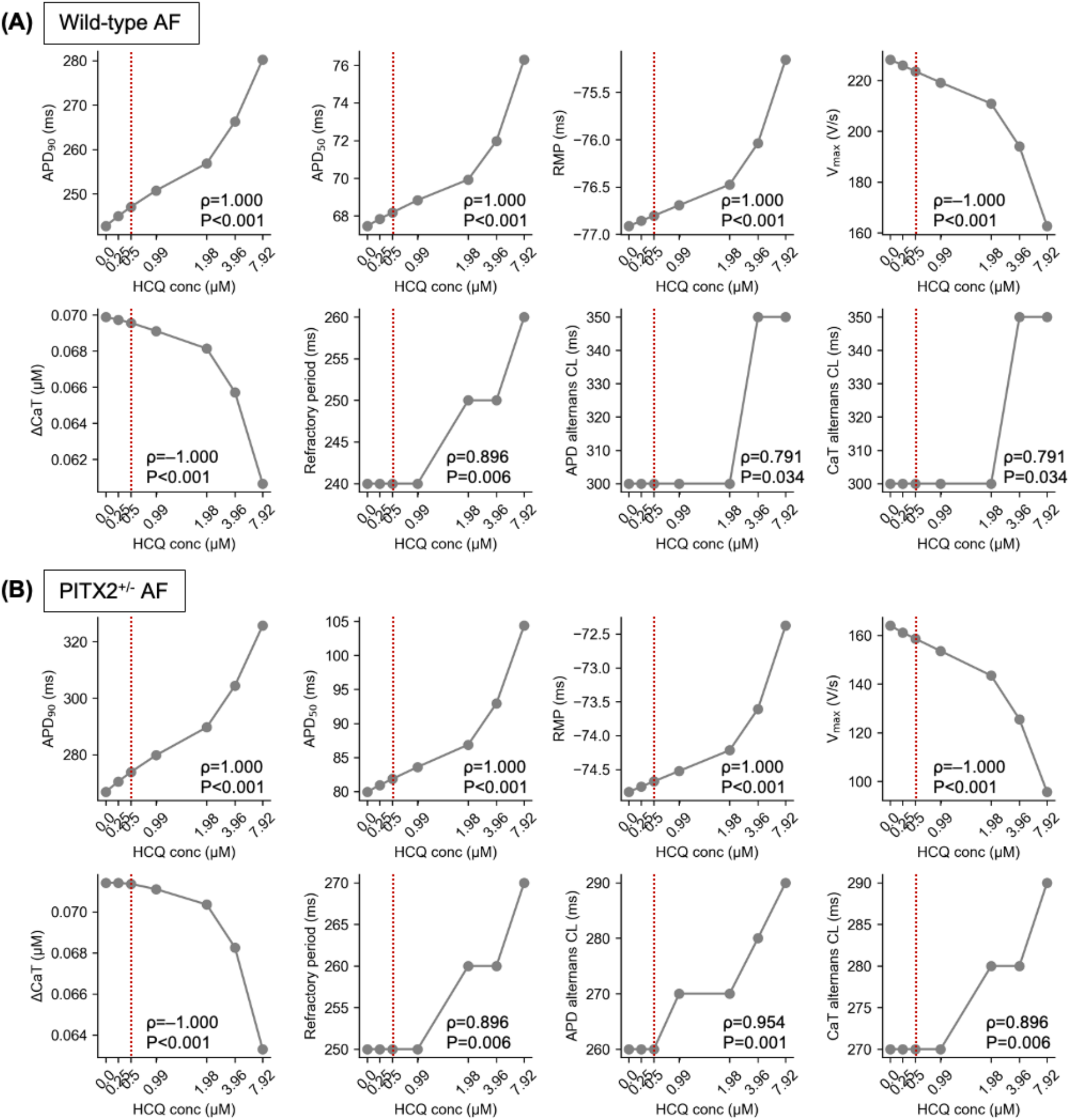
Electrophysiological markers in the wild-type AF (A) and PITX2^+/−^ AF (B) conditions. The action potential duration at 90% of repolarization (APD_90_), APD at 50% of repolarization (APD_50_), resting membrane potential (RMP), maximum upstroke velocity (V_max_), calcium transient amplitude (ΔCaT), refractory period, APD alternans threshold cycle length (CL), and CaT alternans threshold CL are shown, depending on hydroxychloroquine concentrations of 0–16×C_max_ (see Methods). Red lines indicate C_max_. Spearman’s correlation coefficients and P-values are presented. HCQ, hydroxychloroquine; C_max_, maximum free therapeutic plasma concentration; AF, atrial fibrillation.

However, HCQ at 2×C_max_ began to increase the APD alternans threshold in the PITX2^+/−^ AF model (Figure 3B). To examine the effects of HCQ on alternans, we paced *in silico* atrial cells at 4 Hz. As highlighted in Figure 4, HCQ at 4×C_max_ and 16×C_max_ caused prominent alternating long and short action potentials in the PITX2^+/−^ AF condition (APD variation>10%, Figure 4C), whereas the NSR and wild-type AF models did not show prominent APD alternans (APD variation<10%).

**Figure 4.**
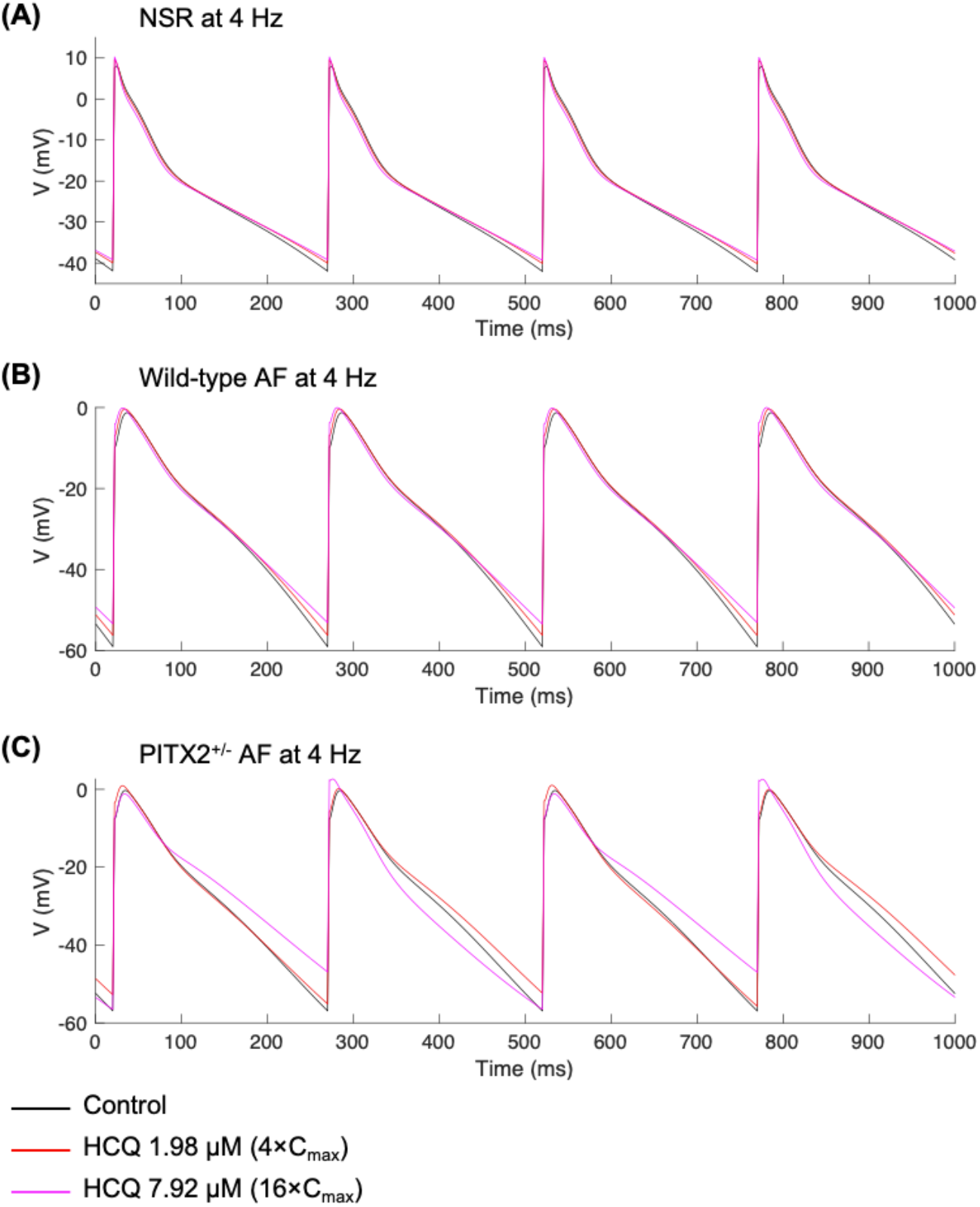
Simulated action potentials (APs) at the pacing frequency of 4 Hz in normal sinus rhythm (A), wild-type AF (B), and PITX2^+/−^ AF (C). With the presence of hydroxychloroquine, alternating long and short APs are prominent in PITX2^+/−^ AF (beat-to-beat APD variation>10%) (C). HCQ, hydroxychloroquine; C_max_, maximum free therapeutic plasma concentration; NSR, normal sinus rhythm; AF, atrial fibrillation.

## 4. Discussion

We systematically measured atrial electrophysiological markers after applying HCQ to *in silico* human atrial cardiomyocytes. We found that HCQ increased the APD and decreased ΔCaT but did not change the alternans thresholds at C_max_. Additionally, HCQ reduced the maximum upstroke velocity, potentially slowing the myocardial conduction velocity. Since AF is characterized by shortening of the APD and rate-dependent APD alternans occurrence [33, 37, 38], our computational results suggest no significant atrial arrhythmogenic effects of HCQ at C_max_, rather indicating a potential antiarrhythmic role of low-dose HCQ in AF. However, with high doses of HCQ and high heart rates, APD alternans were prominent, particularly in the PITX2^+/−^ AF condition. This potential proarrhythmogenic role of high-dose HCQ in PITX2^+/−^ AF should be further investigated experimentally.

Recent studies suggest that HCQ may play a cardioprotective role in patients with autoimmune diseases [39]. Gupta et al. [26] showed that HCQ use was associated with an 88% decrease in the AF risk in patients with SLE. We found that HCQ increased the atrial APD in a dose-dependent manner without affecting the alternans thresholds at C_max_. Our results imply the potential antiarrhythmic effects of HCQ against AF. Other studies have shown that HCQ treatments did not significantly increase the risk of cardiac arrhythmias, including ventricular arrhythmias, in patients with SLE [40, 41] and common rheumatic diseases [42]. Although some case reports showed that HCQ could lead to QT prolongation [13, 43], HCQ use was not associated with QTc length in a large cohort of patients with rheumatic diseases [44]. In contrast, HCQ may increase the risk of life-threatening ventricular arrhythmias in patients with COVID-19 [45]. In patients with COVID-19, HCQ treatment increased the risks of QT interval prolongation, TdP, and cardiac arrest [45–49]. For COVID-19, HCQ is typically started at high dosages of 2000 mg and reduced to 800 mg, which is higher than the typical maintenance dosage of 200 mg in patients with SLE. Therefore, adverse cardiac effects of HCQ in COVID-19 may be attributed to the use of high-dose HCQ, which could lead to prominent APD alternans according to our data (Figure 4). Further large-scale cohort studies are needed to determine the causal effects of HCQ on AF.

Most antiarrhythmic drugs for AF increase the atrial APD. However, an excessively prolonged atrial APD can lead to atrial arrhythmias [50] and concomitant effects on ventricular APD can induce TdP [51, 52]. Our data on atrial electrophysiology and other studies suggest [15, 16] that HCQ increases both the atrial and ventricular APDs. Thus, estimating the safe therapeutic range of HCQ is essential to reduce the risk of AF and TdP. In our study, HCQ increased the atrial APDs by only 1.4–2.6% at C_max_. However, patients with SLE taking daily HCQ doses may have high blood HCQ concentrations due to its long half-life [36]. With high-dose HCQ, we observed prominent APD alternans, particularly in the PITX2^+/−^ AF condition (Figure 4). Therefore, patients with concomitant autoimmune diseases and *PITX2* mutations may have an increased risk of AF, and this potential proarrhythmogenic effect should be further investigated using real-world clinical data.

Besides the direct effects of HCQ on atrial electrophysiology, HCQ may play a cardioprotective role in AF by inhibiting autophagy flux. Yuan et al. [53] discovered that autophagy induces atrial electrical remodeling through ubiquitin-dependent degradation of the L-type calcium channel. Additionally, serum autophagy related 5 (ATG5) was associated with left atrial enlargement in patients with paroxysmal AF [54]. Thus, interruption of autophagy flux by HCQ may prevent the electrical and structural remodeling of AF.

Cardiac computational modeling approaches can be used to quantitatively understand complex subcellular or spatiotemporal dynamics of AF [23, 25, 33] and to perform personalized AF ablation as shown in the OPTIMA [22] and PersonAL [23] studies. Recently, these computational models have also been utilized for *in-silico* antiarrhythmic drug trials and drug toxicity screening [18, 19, 24, 25]. Although we did not perform 2D/3D tissue-level simulations, our computational approaches can be directly extended to examine how HCQ quantitatively affects the spatiotemporal wave dynamics of AF, antiarrhythmic drug efficacies, and ablation outcomes in normal, wild-type AF, and PITX2^+/−^ AF conditions [55]. We also expect that cardiac computational modeling might be helpful for personalizing AF risk stratification and management in patients with SLE taking HCQ.

There are several limitations to our computational study. Although we used a state-of-the-art computational model of the human atrial cardiomyocyte [20], the results need to be validated using human induced pluripotent stem cells, animal models, or clinical electrophysiology data. We assumed fixed baseline parameter conditions; however, population-based approaches would provide more statistically robust results by considering the biological uncertainty/variability in ion channel parameters [17, 25, 31, 32]. Since atrial cell models have pacing protocol-dependent memory effects [56], various pacing protocol-related sensitivity analyses should be considered to obtain more robust and reliable results [57]. To elucidate the comprehensive mechanisms of action of HCQ for drug repurposing in AF, transcriptome/proteome perturbation data or protein-protein interaction network analysis would be helpful [58, 59].

## Acknowledgements

This research received no external funding. This study did not use new animal/clinical data. The author would like to thank anonymous reviewers for their insightful comments.

## Authorship contribution

**Euijun Song:** Conceptualization, Methodology, Software, Investigation, Data curation, Visualization, Writing – original draft, Writing – review & editing.

---

**Supplementary Figure S1.**
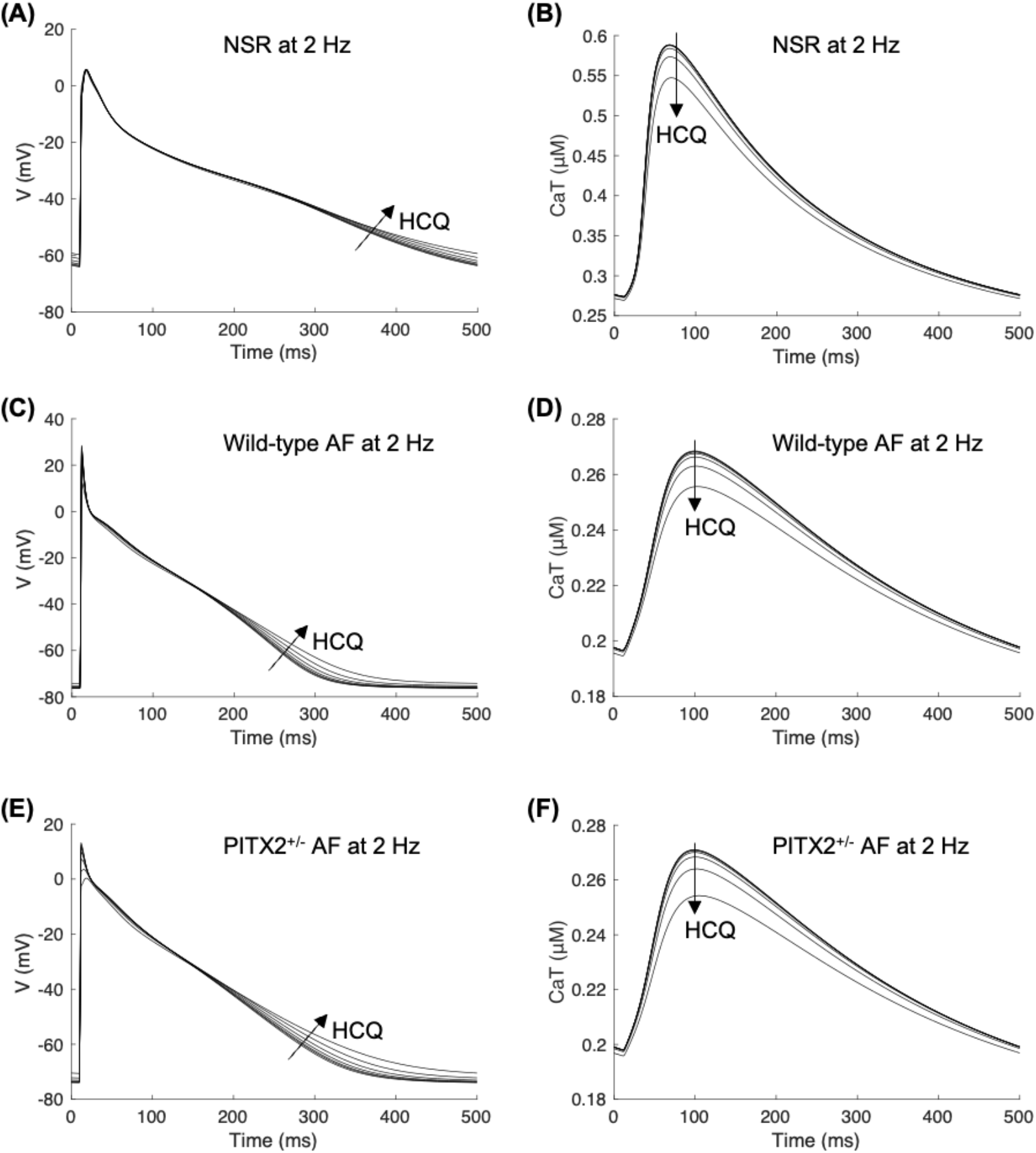
Action potentials (APs) and calcium transients (CaTs) for hydroxychloroquine concentrations of 0–16×C_max_. Simulated steady-state AP and CaT traces are shown at the pacing frequency of 2 Hz in normal sinus rhythm (A, B), wild-type AF (C, D), and PITX2^+/−^ AF (E, F). HCQ, hydroxychloroquine; C_max_, maximum free therapeutic plasma concentration; CaT, calcium transient; NSR, normal sinus rhythm; AF, atrial fibrillation.

## References

[1] E.L. Nirk, F. Reggiori, M. Mauthe, Hydroxychloroquine in rheumatic autoimmune disorders and beyond, EMBO Mol Med, 12 (2020) e12476.

[2] E. Schrezenmeier, T. Dorner, Mechanisms of action of hydroxychloroquine and chloroquine: implications for rheumatology, Nat Rev Rheumatol, 16 (2020) 155–166.

[3] R.C. Group, P. Horby, M. Mafham, L. Linsell, J.L. Bell, N. Staplin, J.R. Emberson, M. Wiselka, A. Ustianowski, E. Elmahi, B. Prudon, T. Whitehouse, T. Felton, J. Williams, J. Faccenda, J. Underwood, J.K. Baillie, L.C. Chappell, S.N. Faust, T. Jaki, K. Jeffery, W.S. Lim, A. Montgomery, K. Rowan, J. Tarning, J.A. Watson, N.J. White, E. Juszczak, R. Haynes, M.J. Landray, Effect of Hydroxychloroquine in Hospitalized Patients with Covid-19, N Engl J Med, 383 (2020) 2030–2040.

[4] D.R. Boulware, M.F. Pullen, A.S. Bangdiwala, K.A. Pastick, S.M. Lofgren, E.C. Okafor, C.P. Skipper, A.A. Nascene, M.R. Nicol, M. Abassi, N.W. Engen, M.P. Cheng, D. LaBar, S.A. Lother, L.J. MacKenzie, G. Drobot, N. Marten, R. Zarychanski, L.E. Kelly, I.S. Schwartz, E.G. McDonald, R. Rajasingham, T.C. Lee, K.H. Hullsiek, A Randomized Trial of Hydroxychloroquine as Postexposure Prophylaxis for Covid-19, N Engl J Med, 383 (2020) 517–525.

[5] A.B. Cavalcanti, F.G. Zampieri, R.G. Rosa, L.C.P. Azevedo, V.C. Veiga, A. Avezum, L.P. Damiani, A. Marcadenti, L. Kawano-Dourado, T. Lisboa, D.L.M. Junqueira, E.S.P.G.M. de Barros, L. Tramujas, E.O. Abreu-Silva, L.N. Laranjeira, A.T. Soares, L.S. Echenique, A.J. Pereira, F.G.R. Freitas, O.C.E. Gebara, V.C.S. Dantas, R.H.M. Furtado, E.P. Milan, N.A. Golin, F.F. Cardoso, I.S. Maia, C.R. Hoffmann Filho, A.P.M. Kormann, R.B. Amazonas, M.F. Bocchi de Oliveira, A. Serpa-Neto, M. Falavigna, R.D. Lopes, F.R. Machado, O. Berwanger, I.I. Coalition Covid-19 Brazil, Hydroxychloroquine with or without Azithromycin in Mild-to-Moderate Covid-19, N Engl J Med, 383 (2020) 2041–2052.

[6] K. Sacre, L.A. Criswell, J.M. McCune, Hydroxychloroquine is associated with impaired interferon-alpha and tumor necrosis factor-alpha production by plasmacytoid dendritic cells in systemic lupus erythematosus, Arthritis Res Ther, 14 (2012) R155.

[7] D.J. Wallace, M. Linker-Israeli, A.L. Metzger, V.J. Stecher, The relevance of antimalarial therapy with regard to thrombosis, hypercholesterolemia and cytokines in SLE, Lupus, 2 Suppl 1 (1993) S13–15.

[8] R. Banchereau, S. Hong, B. Cantarel, N. Baldwin, J. Baisch, M. Edens, A.M. Cepika, P. Acs, J. Turner, E. Anguiano, P. Vinod, S. Kahn, G. Obermoser, D. Blankenship, E. Wakeland, L. Nassi, A. Gotte, M. Punaro, Y.J. Liu, J. Banchereau, J. Rossello-Urgell, T. Wright, V. Pascual, Personalized Immunomonitoring Uncovers Molecular Networks that Stratify Lupus Patients, Cell, 165 (2016) 551–565.

[9] C. Ponticelli, G. Moroni, Hydroxychloroquine in systemic lupus erythematosus (SLE), Expert Opin Drug Saf, 16 (2017) 411–419.

[10] C. Chatre, F. Roubille, H. Vernhet, C. Jorgensen, Y.M. Pers, Cardiac Complications Attributed to Chloroquine and Hydroxychloroquine: A Systematic Review of the Literature, Drug Saf, 41 (2018) 919–931.

[11] N. Costedoat-Chalumeau, J.S. Hulot, Z. Amoura, G. Leroux, P. Lechat, C. Funck-Brentano, J.C. Piette, Heart conduction disorders related to antimalarials toxicity: an analysis of electrocardiograms in 85 patients treated with hydroxychloroquine for connective tissue diseases, Rheumatology (Oxford), 46 (2007) 808–810.

[12] A. Jorge, C. Ung, L.H. Young, R.B. Melles, H.K. Choi, Hydroxychloroquine retinopathy - implications of research advances for rheumatology care, Nat Rev Rheumatol, 14 (2018) 693–703.

[13] J.P. O’Laughlin, P.H. Mehta, B.C. Wong, Life Threatening Severe QTc Prolongation in Patient with Systemic Lupus Erythematosus due to Hydroxychloroquine, Case Rep Cardiol, 2016 (2016) 4626279.

[14] L. Jankelson, G. Karam, M.L. Becker, L.A. Chinitz, M.C. Tsai, QT prolongation, torsades de pointes, and sudden death with short courses of chloroquine or hydroxychloroquine as used in COVID-19: A systematic review, Heart Rhythm, 17 (2020) 1472–1479.

[15] U. Thomet, B. Amuzescu, T. Knott, S.A. Mann, K. Mubagwa, B.M. Radu, Assessment of proarrhythmogenic risk for chloroquine and hydroxychloroquine using the CiPA concept, European Journal of Pharmacology, 913 (2021) 174632.

[16] A. Delaunois, M. Abernathy, W.D. Anderson, K.A. Beattie, K.W. Chaudhary, J. Coulot, V. Gryshkova, S. Hebeisen, M. Holbrook, J. Kramer, Y. Kuryshev, D. Leishman, I. Lushbough, E. Passini, W.S. Redfern, B. Rodriguez, E.I. Rossman, C. Trovato, C. Wu, J.P. Valentin, Applying the CiPA approach to evaluate cardiac proarrhythmia risk of some antimalarials used off-label in the first wave of COVID-19, Clin Transl Sci, 14 (2021) 1133–1146.

[17] D.G. Whittaker, R.A. Capel, M. Hendrix, X.H.S. Chan, N. Herring, N.J. White, G.R. Mirams, R.B. Burton, Cardiac TdP risk stratification modelling of anti-infective compounds including chloroquine and hydroxychloroquine, R Soc Open Sci, 8 (2021) 210235.

[18] Z. Li, B.J. Ridder, X. Han, W.W. Wu, J. Sheng, P.N. Tran, M. Wu, A. Randolph, R.H. Johnstone, G.R. Mirams, Y. Kuryshev, J. Kramer, C. Wu, W.J. Crumb, Jr., D.G. Strauss, Assessment of an In Silico Mechanistic Model for Proarrhythmia Risk Prediction Under the CiPA Initiative, Clin Pharmacol Ther, 105 (2019) 466–475.

[19] Z. Li, S. Dutta, J. Sheng, P.N. Tran, W. Wu, K. Chang, T. Mdluli, D.G. Strauss, T. Colatsky, Improving the In Silico Assessment of Proarrhythmia Risk by Combining hERG (Human Ether-a-go-go-Related Gene) Channel-Drug Binding Kinetics and Multichannel Pharmacology, Circ Arrhythm Electrophysiol, 10 (2017) e004628.

[20] E. Grandi, S.V. Pandit, N. Voigt, A.J. Workman, D. Dobrev, J. Jalife, D.M. Bers, Human atrial action potential and Ca2+ model: sinus rhythm and chronic atrial fibrillation, Circ Res, 109 (2011) 1055–1066.

[21] M. Courtemanche, R.J. Ramirez, S. Nattel, Ionic mechanisms underlying human atrial action potential properties: insights from a mathematical model, Am J Physiol, 275 (1998) H301–321.

[22] P.M. Boyle, T. Zghaib, S. Zahid, R.L. Ali, D. Deng, W.H. Franceschi, J.B. Hakim, M.J. Murphy, A. Prakosa, S.L. Zimmerman, H. Ashikaga, J.E. Marine, A. Kolandaivelu, S. Nazarian, D.D. Spragg, H. Calkins, N.A. Trayanova, Computationally guided personalized targeted ablation of persistent atrial fibrillation, Nat Biomed Eng, 3 (2019) 870–879.

[23] L. Azzolin, M. Eichenlaub, C. Nagel, D. Nairn, J. Sanchez, L. Unger, O. Dossel, A. Jadidi, A. Loewe, Personalized ablation vs. conventional ablation strategies to terminate atrial fibrillation and prevent recurrence, Europace, (2022).

[24] J.D. Moreno, Z.I. Zhu, P.C. Yang, J.R. Bankston, M.T. Jeng, C. Kang, L. Wang, J.D. Bayer, D.J. Christini, N.A. Trayanova, C.M. Ripplinger, R.S. Kass, C.E. Clancy, A computational model to predict the effects of class I anti-arrhythmic drugs on ventricular rhythms, Sci Transl Med, 3 (2011) 98ra83.

[25] A. Dasi, A. Roy, R. Sachetto, J. Camps, A. Bueno-Orovio, B. Rodriguez, In-silico drug trials for precision medicine in atrial fibrillation: From ionic mechanisms to electrocardiogram-based predictions in structurally-healthy human atria, Front Physiol, 13 (2022) 966046.

[26] A. Gupta, K.J. Shields, S. Manzi, M.C. Wasko, T.S. Sharma, Association of Hydroxychloroquine Use With Decreased Incident Atrial Fibrillation in Systemic Lupus Erythematosus, Arthritis Care Res (Hoboken), 73 (2021) 828–832.

[27] F. Syeda, A.P. Holmes, T.Y. Yu, S. Tull, S.M. Kuhlmann, D. Pavlovic, D. Betney, G. Riley, J.P. Kucera, F. Jousset, J.R. de Groot, S. Rohr, N.A. Brown, L. Fabritz, P. Kirchhof, PITX2 Modulates Atrial Membrane Potential and the Antiarrhythmic Effects of Sodium-Channel Blockers, J Am Coll Cardiol, 68 (2016) 1881–1894.

[28] D.F. Gudbjartsson, D.O. Arnar, A. Helgadottir, S. Gretarsdottir, H. Holm, A. Sigurdsson, A. Jonasdottir, A. Baker, G. Thorleifsson, K. Kristjansson, A. Palsson, T. Blondal, P. Sulem, V.M. Backman, G.A. Hardarson, E. Palsdottir, A. Helgason, R. Sigurjonsdottir, J.T. Sverrisson, K. Kostulas, M.C. Ng, L. Baum, W.Y. So, K.S. Wong, J.C. Chan, K.L. Furie, S.M. Greenberg, M. Sale, P. Kelly, C.A. MacRae, E.E. Smith, J. Rosand, J. Hillert, R.C. Ma, P.T. Ellinor, G. Thorgeirsson, J.R. Gulcher, A. Kong, U. Thorsteinsdottir, K. Stefansson, Variants conferring risk of atrial fibrillation on chromosome 4q25, Nature, 448 (2007) 353–357.

[29] S.A. Lubitz, M.F. Sinner, K.L. Lunetta, S. Makino, A. Pfeufer, R. Rahman, C.E. Veltman, J. Barnard, J.C. Bis, S.P. Danik, A. Sonni, M.A. Shea, F. Del Monte, S. Perz, M. Muller, A. Peters, S.M. Greenberg, K.L. Furie, C. van Noord, E. Boerwinkle, B.H. Stricker, J. Witteman, J.D. Smith, M.K. Chung, S.R. Heckbert, E.J. Benjamin, J. Rosand, D.E. Arking, A. Alonso, S. Kaab, P.T. Ellinor, Independent susceptibility markers for atrial fibrillation on chromosome 4q25, Circulation, 122 (2010) 976–984.

[30] F. Syeda, P. Kirchhof, L. Fabritz, PITX2-dependent gene regulation in atrial fibrillation and rhythm control, J Physiol, 595 (2017) 4019–4026.

[31] M.R. Vagos, H. Arevalo, B.L. de Oliveira, J. Sundnes, M.M. Maleckar, A computational framework for testing arrhythmia marker sensitivities to model parameters in functionally calibrated populations of atrial cells, Chaos, 27 (2017) 093941.

[32] E. Song, Y.S. Lee, Interpretable machine learning of action potential duration restitution kinetics in single-cell models of atrial cardiomyocytes, J Electrocardiol, 74 (2022) 137–145.

[33] Y.S. Lee, M. Hwang, J.S. Song, C. Li, B. Joung, E.A. Sobie, H.N. Pak, The Contribution of Ionic Currents to Rate-Dependent Action Potential Duration and Pattern of Reentry in a Mathematical Model of Human Atrial Fibrillation, PLoS One, 11 (2016) e0150779.

[34] G.R. Mirams, Y. Cui, A. Sher, M. Fink, J. Cooper, B.M. Heath, N.C. McMahon, D.J. Gavaghan, D. Noble, Simulation of multiple ion channel block provides improved early prediction of compounds’ clinical torsadogenic risk, Cardiovasc Res, 91 (2011) 53–61.

[35] X. Yao, F. Ye, M. Zhang, C. Cui, B. Huang, P. Niu, X. Liu, L. Zhao, E. Dong, C. Song, S. Zhan, R. Lu, H. Li, W. Tan, D. Liu, In Vitro Antiviral Activity and Projection of Optimized Dosing Design of Hydroxychloroquine for the Treatment of Severe Acute Respiratory Syndrome Coronavirus 2 (SARS-CoV-2), Clin Infect Dis, 71 (2020) 732–739.

[36] N. Costedoat-Chalumeau, Z. Amoura, J.S. Hulot, H.A. Hammoud, G. Aymard, P. Cacoub, C. Frances, B. Wechsler, L.T. Huong du, P. Ghillani, L. Musset, P. Lechat, J.C. Piette, Low blood concentration of hydroxychloroquine is a marker for and predictor of disease exacerbations in patients with systemic lupus erythematosus, Arthritis Rheum, 54 (2006) 3284–3290.

[37] t.J. Wu, Y.H. Kim, M. Yashima, C.A. Athill, C.T. Ting, H.S. Karagueuzian, P.S. Chen, Progressive action potential duration shortening and the conversion from atrial flutter to atrial fibrillation in the isolated canine right atrium, J Am Coll Cardiol, 38 (2001) 1757–1765.

[38] M.R. Franz, S.M. Jamal, S.M. Narayan, The role of action potential alternans in the initiation of atrial fibrillation in humans: a review and future directions, Europace, 14 Suppl 5 (2012) v58–v64.

[39] C.C. Prodromos, T. Rumschlag, T. Perchyk, Hydroxychloroquine is protective to the heart, not harmful: a systematic review, New Microbes New Infect, 37 (2020) 100747.

[40] C.H. Lo, Y.H. Wang, C.F. Tsai, K.C. Chan, L.C. Li, T.H. Lo, J.C. Wei, C.H. Su, Association of hydroxychloroquine and cardiac arrhythmia in patients with systemic lupus erythematosus: A population-based case control study, PLoS One, 16 (2021) e0251918.

[41] t.K. McGhie, P. Harvey, J. Su, N. Anderson, G. Tomlinson, Z. Touma, Electrocardiogram abnormalities related to anti-malarials in systemic lupus erythematosus, Clin Exp Rheumatol, 36 (2018) 545–551.

[42] C.H. Lo, J.C. Wei, Y.H. Wang, C.F. Tsai, K.C. Chan, L.C. Li, T.H. Lo, C.H. Su, Hydroxychloroquine Does Not Increase the Risk of Cardiac Arrhythmia in Common Rheumatic Diseases: A Nationwide Population-Based Cohort Study, Front Immunol, 12 (2021) 631869.

[43] N.D. Morgan, S.V. Patel, O. Dvorkina, Suspected hydroxychloroquine-associated QT-interval prolongation in a patient with systemic lupus erythematosus, J Clin Rheumatol, 19 (2013) 286–288.

[44] E. Park, J.T. Giles, T. Perez-Recio, P. Pina, C. Depender, Y. Gartshteyn, A.D. Askanase, J. Bathon, L. Geraldino-Pardilla, Hydroxychloroquine use is not associated with QTc length in a large cohort of SLE and RA patients, Arthritis Res Ther, 23 (2021) 271.

[45] I.M. Tleyjeh, Z. Kashour, O. AlDosary, M. Riaz, H. Tlayjeh, M.A. Garbati, R. Tleyjeh, M.H. Al-Mallah, M.R. Sohail, D. Gerberi, A.A. Bin Abdulhak, J.R. Giudicessi, M.J. Ackerman, T. Kashour, Cardiac Toxicity of Chloroquine or Hydroxychloroquine in Patients With COVID-19: A Systematic Review and Meta-regression Analysis, Mayo Clin Proc Innov Qual Outcomes, 5 (2021) 137–150.

[46] M. Saleh, J. Gabriels, D. Chang, B. Soo Kim, A. Mansoor, E. Mahmood, P. Makker, H. Ismail, B. Goldner, J. Willner, S. Beldner, R. Mitra, R. John, J. Chinitz, N. Skipitaris, S. Mountantonakis, L.M. Epstein, Effect of Chloroquine, Hydroxychloroquine, and Azithromycin on the Corrected QT Interval in Patients With SARS-CoV-2 Infection, Circ Arrhythm Electrophysiol, 13 (2020) e008662.

[47] E. Chorin, L. Wadhwani, S. Magnani, M. Dai, E. Shulman, C. Nadeau-Routhier, R. Knotts, R. Bar-Cohen, E. Kogan, C. Barbhaiya, A. Aizer, D. Holmes, S. Bernstein, M. Spinelli, D.S. Park, C. Stefano, L.A. Chinitz, L. Jankelson, QT interval prolongation and torsade de pointes in patients with COVID-19 treated with hydroxychloroquine/azithromycin, Heart Rhythm, 17 (2020) 1425–1433.

[48] N.J. Mercuro, C.F. Yen, D.J. Shim, T.R. Maher, C.M. McCoy, P.J. Zimetbaum, H.S. Gold, Risk of QT Interval Prolongation Associated With Use of Hydroxychloroquine With or Without Concomitant Azithromycin Among Hospitalized Patients Testing Positive for Coronavirus Disease 2019 (COVID-19), JAMA Cardiol, 5 (2020) 1036–1041.

[49] E.S. Rosenberg, E.M. Dufort, T. Udo, L.A. Wilberschied, J. Kumar, J. Tesoriero, P. Weinberg, J. Kirkwood, A. Muse, J. DeHovitz, D.S. Blog, B. Hutton, D.R. Holtgrave, H.A. Zucker, Association of Treatment With Hydroxychloroquine or Azithromycin With In-Hospital Mortality in Patients With COVID-19 in New York State, JAMA, 323 (2020) 2493–2502.

[50] P. Kirchhof, L. Eckardt, M.R. Franz, G. Monnig, P. Loh, H. Wedekind, E. Schulze-Bahr, G. Breithardt, W. Haverkamp, Prolonged atrial action potential durations and polymorphic atrial tachyarrhythmias in patients with long QT syndrome, J Cardiovasc Electrophysiol, 14 (2003) 1027–1033.

[51] J. Shenthar, J.M. Rachaiah, V. Pillai, S.S. Chakali, V. Balasubramanian, M. Chollenhalli Nanjappa, Incidence of drug-induced torsades de pointes with intravenous amiodarone, Indian Heart J, 69 (2017) 707–713.

[52] H.E. Lim, H.N. Pak, J.C. Ahn, W.H. Song, Y.H. Kim, Torsade de pointes induced by short-term oral amiodarone therapy, Europace, 8 (2006) 1051–1053.

[53] Y. Yuan, J. Zhao, Y. Gong, D. Wang, X. Wang, F. Yun, Z. Liu, S. Zhang, W. Li, X. Zhao, L. Sun, L. Sheng, Z. Pan, Y. Li, Autophagy exacerbates electrical remodeling in atrial fibrillation by ubiquitin-dependent degradation of L-type calcium channel, Cell Death Dis, 9 (2018) 873.

[54] H. Fedai, I.H. Altiparmak, M.B. Tascanov, Z. Tanriverdi, A. Bicer, F. Gungoren, R. Demirbag, I. Koyuncu, The relationship between oxidative stress and autophagy and apoptosis in patients with paroxysmal atrial fibrillation, Scand J Clin Lab Invest, 82 (2022) 391–397.

[55] J. Bai, A. Lo, P.A. Gladding, M.K. Stiles, V.V. Fedorov, J. Zhao, In silico investigation of the mechanisms underlying atrial fibrillation due to impaired Pitx2, PLoS Comput Biol, 16 (2020) e1007678.

[56] E.M. Cherry, H.M. Hastings, S.J. Evans, Dynamics of human atrial cell models: restitution, memory, and intracellular calcium dynamics in single cells, Prog Biophys Mol Biol, 98 (2008) 24–37.

[57] L. Azzolin, S. Schuler, O. Dossel, A. Loewe, A Reproducible Protocol to Assess Arrhythmia Vulnerability in silico: Pacing at the End of the Effective Refractory Period, Front Physiol, 12 (2021) 656411.

[58] E. Song, R.S. Wang, J.A. Leopold, J. Loscalzo, Network determinants of cardiovascular calcification and repositioned drug treatments, FASEB J, 34 (2020) 11087–11100.

[59] J.C. Lal, C. Mao, Y. Zhou, S.R. Gore-Panter, J.H. Rennison, B.S. Lovano, L. Castel, J. Shin, A.M. Gillinov, J.D. Smith, J. Barnard, D.R. Van Wagoner, Y. Luo, F. Cheng, M.K. Chung, Transcriptomics-based network medicine approach identifies metformin as a repurposable drug for atrial fibrillation, Cell Rep Med, 3 (2022) 100749.

